## Supplementary figures and images for "NIRis: A low-cost, versatile imaging system for NIR fluorescence detection of phototrophic cell colonies used in science and education"

### S1 Fig: Individual 3D-printed parts

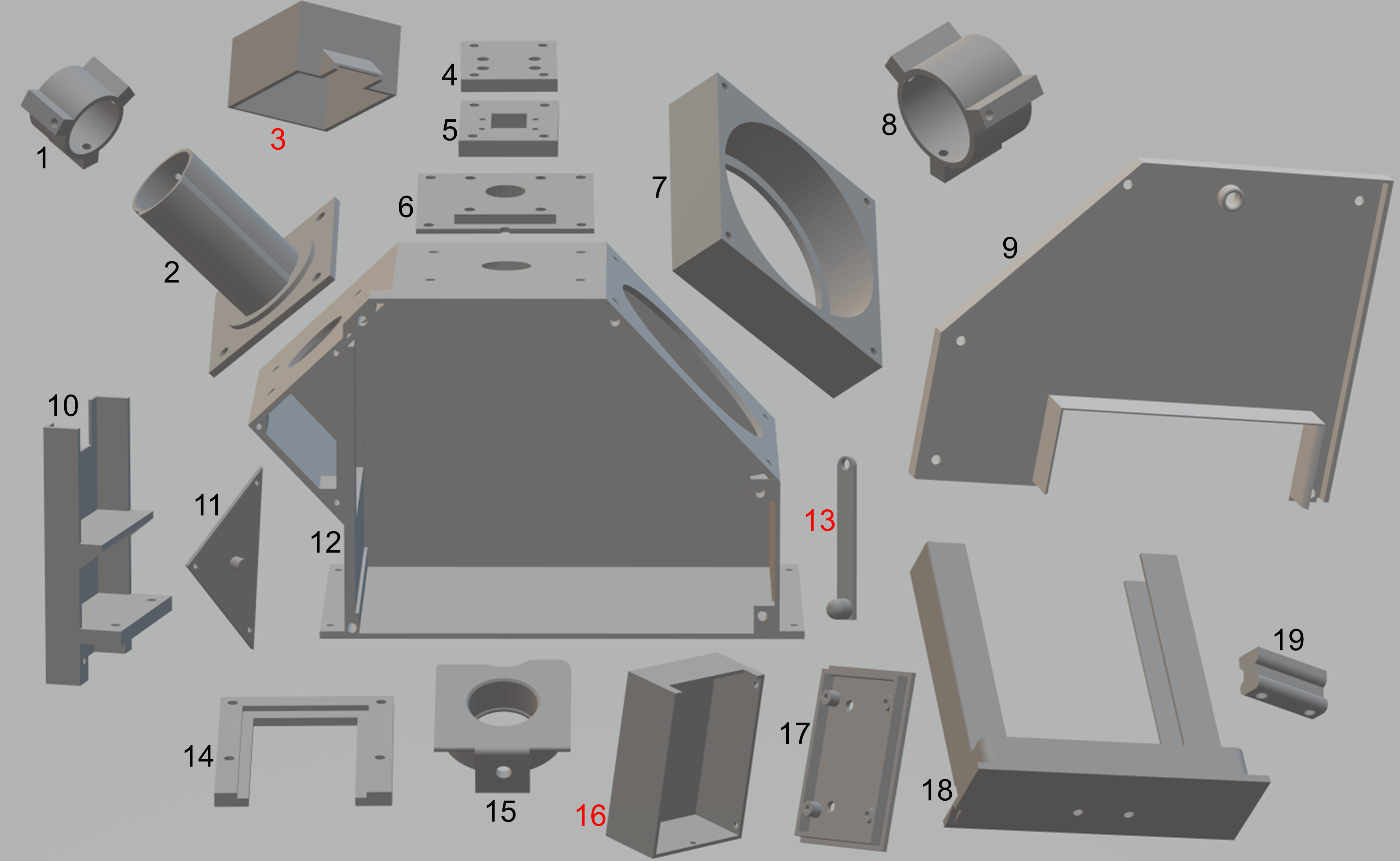

### S2 Fig: Fluorescence spectra of a typical AAPB

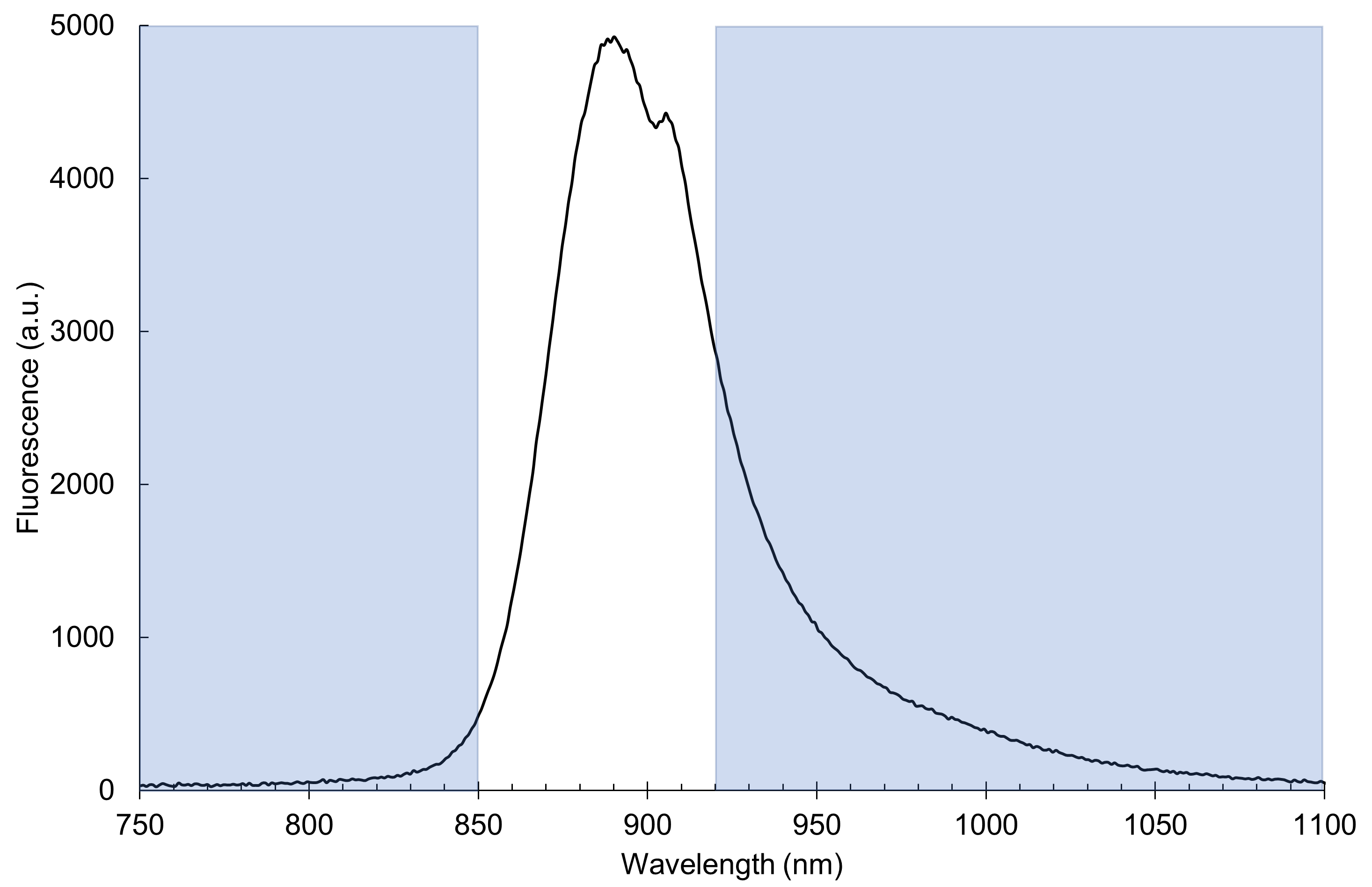
